# Sequence representations and their utility for predicting protein-protein interactions

**DOI:** 10.1101/2019.12.31.890699

**Authors:** Dhananjay Kimothi, Pravesh Biyani, James M Hogan

## Abstract

Protein-Protein Interactions (PPIs) are a crucial mechanism underpinning the function of the cell. Predicting the likely relationship between a pair of proteins is thus an important problem in bioinformatics, and a wide range of machine-learning based methods have been proposed for this task. Their success is heavily dependent on the construction of the feature vectors, with most using a set of physico-chemical properties derived from the sequence. Few work directly with the sequence itself.

Recent works on embedding sequences in a low dimensional vector space has shown the utility of this approach for tasks such as protein classification and sequence search. In this paper, we extend these ideas to the PPI task, making inferences from the pair instead of for the individual sequences. We evaluate the method on human and yeast PPI datasets, benchmarking against the established methods. These results demonstrate that we can obtain sequence encodings for the PPI task which achieve similar levels of performance to existing methods without reliance on complex physico-chemical feature sets.

## 1 Introduction

Proteins play a central role in cellular function, often mediated through the protein interactions leading to the formation of transient or stable complexes. Such Protein-Protein interactions (PPIs) vary in their duration and character, being governed principally by electrostatic forces and constrained by molecular structure [1]. Knowledge of PPIs can help determine the function of new proteins and improve our understanding of cellular processes such as DNA transcription and metabolic cycles [2].

A number of experimental techniques have been developed for determining the nature and extent of interactions between proteins, and approaches such as the yeast two-hybrid system [3], mass spectrometry [4] and protein chips [5] have been widely used. Yet experimental methods share two significant drawbacks: (i) such methods are expensive and time-consuming, and (ii) they are known to exhibit high false-positive rates [6]. To counter these limitations, researchers have developed alternative computational methods [7–14] for predicting PPIs. These methods rely on the extraction and processing of features from biological sequences and their associated metadata. The earliest work in this direction was based on identifying domains [7–9], sequence conservation between interacting proteins [10,11] and exploiting structural information [12–14].

The subsequent works focused on machine learning approaches, the success of which primarily rely on representing protein sequences as a *n*-dimensional feature vector. These feature vectors essentially summarize specific attributes of a protein sequence – such as the distribution of amino acids, their physicochemical properties [15], or the composition of a localized region [16,17]. Since for PPI prediction problem, the input is protein pairs, to apply machine learning approaches directly over them, for each pair, we first need to obtain its joint representation. Generally, such representation for a protein pair is obtained by concatenating the feature vector of its constituent protein sequences, hence keeping the information derived from them intact. Although the sequence encoding methods discussed above are predominantly used by the researchers for PPI prediction problem, they suffer from three significant drawbacks: (i) such methods require prior knowledge of physico-chemical properties, (ii) the properties used are not guaranteed to cover the PPI interaction information adequately and (iii) the resulting feature vectors constructed are usually of high dimension, increasing training time and potentially limiting the effectiveness of the model.

One potential solution to these problems is to learn sequence embeddings that capture biological information that may be present in the sequences itself and use them for representing a protein pair. The earliest work on learning biological sequence embeddings by Asgari et.al [18] shows that it is possible to learn biological-information rich embeddings for sun-sequences (*kmers*) directly using the sequence data. They and subsequently Kimothi et al. [19] also demonstrated the utility of such learning techniques, generally called as representation learning (RepL) for the protein classification task. Following these primary works, many of the research has shown the effectiveness of RepL approaches for a range of other downstream bioinformatics tasks [20–23] such as trans-membrane and localization prediction. Although learning of sequence embeddings and their usages for downstream bioinformatics tasks is been explored now for last few years, but to date, such methods have not been studied in detail for PPI prediction problem.

In this paper, we propose to use sequence embeddings learned through RepL methods to represent a protein pair for making PPI predictions. Similar to the other established methods, to represent a protein pair, we concatenate the embedding of its constituent sequences. Here, we hypothesize that the relationship between two proteins can be better determined using the contextual information-rich low dimensional sequence embeddings as compared to the high dimensional physicochemical properties based sequence feature vectors. Note that RepL methods encode the contextual (local) information present in the sequences to generate the corresponding low dimensional embedding.

Apart from the new representation for protein pairs, in this paper, we also focus on the extensive evaluation and comparison with other protein interaction prediction methods. Most of the established methods, including the most recent, use a standard cross-validation setup for evaluation. In this setup, the data is split into training and test set considering the protein pairs but not their constituent proteins. In such a scenario, many proteins (although in different samples) may be present both in the training and test set. In consequence, the prediction of test samples is affected based on the presence or absence of their constituent proteins in the training set. Hence, as noted by Park et al. [24] such an assessment may not be reflective of the ability of the evaluated methods to generalize for paired input.

Given these findings, we rigorously evaluate our proposed approach according to the evaluation strategy proposed by Park et al. [24]. Here the test sets are based on the presence or absence of the individual protein (from the testing sample) in the training set and hence cover different possible scenarios. For experiments, we use benchmark *Human* and *Yeast* datasets provided in [24] and [15]. The experimental results on these datasets show that even the vanilla embedding approaches are capable of achieving better results than most of the physicochemical properties based vector representation. When compared to the recent deep neural network-based approaches applied to sequence data, we find that even a simple framework like ours can give comparable performance stressing the need to better understand the relationship between sequences and the possibility of their interaction. In summary, following are the main contribution of this paper:

- We present a generic PPI prediction framework that uses RepL approaches for feature construction.
- We show that the low dimensional representations learned from the sequences provide better results than most of the feature vectors that exploit physicochemical properties on the PPI prediction task.
- We show that our approach gives comparable performance when compared to much complex end-to-end deep learning classification models.

Before discussing the details of the proposed framework, we briefly review the feature construction methods commonly used for the PPI prediction problem.

### 1.1 Previous Work

The most prominent feature extraction methods include Auto Covariance (AC), Auto Cross Covariance (ACC), Conjoint Triad (CT) and Local Protein Descriptors (LD). AC and ACC based feature vectors are derived by considering protein sequences as time series and computing their auto and cross-correlation properties. The length of the AC and ACC based feature vectors are 420 and 2940 respectively. The conjoint triad method divides the physico-chemical properties of the system into seven categories based on their dipole and volume scale [25], allowing the representation of the protein sequence based on an array of category labels. Once converted, the feature vector of the sequence is given as the frequency of conjoint triads i.e. the three category labels. For a protein pair, the CT method gives a 686-dimensional feature vector.

The Local Descriptor method uses the same categories of physico-chemical properties as CT. The feature vector is constructed from three descriptors – composition (C), Transition (T) and distribution (D) for local regions. The descriptors calculated for each of the local regions are then stacked together to give a 630-dimensional vector for each protein sequence. Other methods based on local regions include Multi-scale Local Feature descriptors (MLD) [26] and the Multi-scale Continuous and Discontinuous feature set (MCD) [27]. MLD uses the multi-scale decomposition of protein sequences to account for overlapping local regions, binary coding to produce an 1134-dimensional vector for a protein pair. MCD works on a similar principle as MLD, though it gives 1764-dimensional vector.

Apart from these methods, approaches such as the physico-chemical Property Response Matrix combined with Local Phase Quantization descriptor (PR-LPQ) [28] and Substitution Matrix Representation (SMR) [29] use signal and image processing techniques to compute the feature vector for a sequence. These methods generally operate on an intermediary matrix representation of a sequence; PR-LPQ uses physico-chemical properties of amino acids, whereas SMR uses the BLOSUM62 [30] matrix to construct the representations.

On top of these feature vectors, a number of different machine learning approaches have been applied to characterise PPIs. These methods have included the Support Vector Machine [15], Random Forests [29] and autoencoders [6]. More recently, studies such as DPPI [31], DNN-PPI [32], and PIPR [33] have explored deep learning frameworks for PPI prediction. Note that these newer deep learning-based approaches are end-to-end classification models and do not specifically focus on the feature construction technique. Also, given their deep architecture, these are complex and hence computationally expensive approaches. Since the success of PPI prediction methods is strongly dependent on the feature vectors extracted from the sequences, new methods for feature extraction remain an active area for researchers.

In the sections below we discuss the proposed framework in detail followed by experiments and results.

## 2 PPI prediction and Proposed Framework

### Notations

We denote *M* protein pairs as 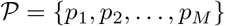, where *p_z_* consist of two samples from a corpus of *N* protein sequences, 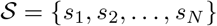. We write *p_z_* = (*s_i_, S_j_*), to uniquely map the sequences with samples in 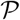, we add superscript to *s_i_* i.e., *p_z_* is written as 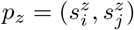. We denote the vocabulary of M unique *k-mers* created from *S* as 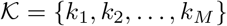 and each sequence *s_i_* = [*k*_*i*_1__, *k*_*i*_2__,…, *k_i_n_i___*] as an ordered list of *k-mers*. To avoid notation clutter, we use *s_i_* also to denote its tag. Finally, the embeddings of *k-mer k_i_*, sequence *s_i_* and protein pair *p_z_* are denoted as **k**_*i*_, **s**_*i*_ and **p**_*z*_ respectively.

### 2.1 PPI prediction problem

PPI prediction is a paired input problem in which the prediction about the relationship between two objects (here proteins) is made. Since machine learning approaches generally operate on a single mathematical entity, for paired input problems, the pair is often represented as a combination of the feature vectors for the individual objects. In a setting where 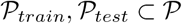 are the training and test set of protein pairs respectively and 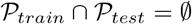, the task in PPI prediction problem is to determine if the proteins in 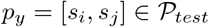 interacts or not given that the label for samples in training set is available.

In our proposed framework, we use RepL methods, like `BioVec, Seq2Vec, SuperVecX` for generating the sequence embedding that are concatenated for a protein pair, and a binary classifier for making predictions. For the better exposition of the proposed framework, in the section below, we first give an overview of the recent biological sequence representation learning (RepL) methods followed by a description of the complete pipeline of the proposed framework, as shown in Fig 2.

**Fig 1.**
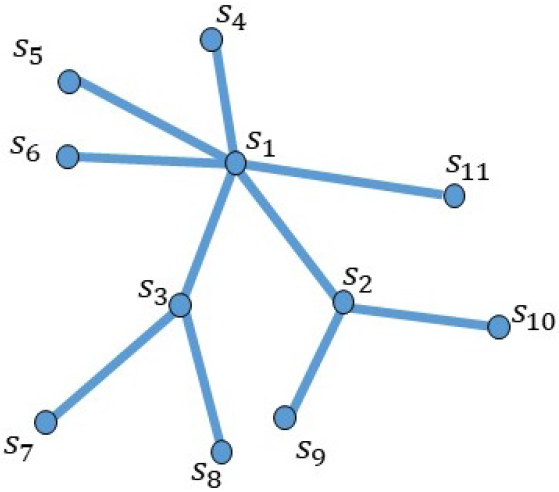
An example of a PPI network obtained from the training pair. Here *s_i_* for *i* = 1, 2, …11 represents the tag of the protein sequences. An edge between two proteins indicate that they interact, missing edge indicates the absence of known interactions.

**Fig 2.**
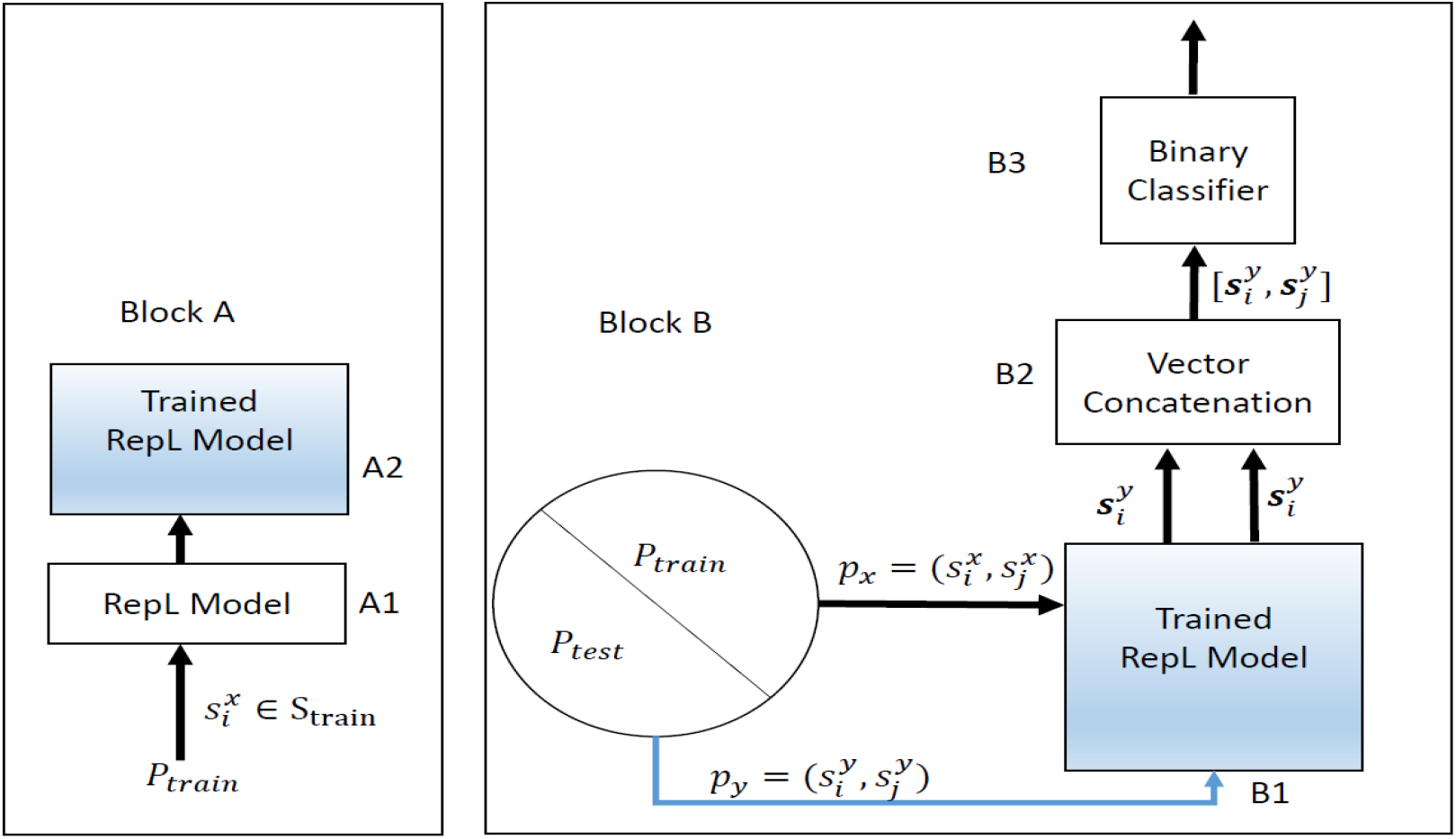
Proposed framework: Block A – training stage of representation learning model; Block B – constitutes a) Train/Test split of interacting/non interacting pair of protein sequences, b) vector concatenation module – for a pair of protein from Train/Test set, the vectors obtained from RL block are concatenated before finally passing to the c) binary classifier

### 2.2 Representation learning methods for biological sequences

Advances in word embedding methods in text processing have inspired several studies of sequence encoding in the bioinformatics domain. These encoding schemes treat sequences as ‘sentences’ and the sub-sequences as ‘words.’ Since the sequences are a continuous array of alphabets and there is no notion of ‘words,’ they are first split by passing a window of size *k* through them. The sub-sequences *(k-mers*) thus generated, are treated as words.

Some of the earliest biological sequence embedding methods include `BioVec` [18] and `Seq2Vec` [34]. Both of these papers utilize `Word2Vec` [19] based architecture for computing the vector representation for protein sequences and demonstrated the utility of such representations for protein family classification tasks. These models are shallow neural network with one hidden layer where the weight matrix between the input/output and hidden layer constitutes the *n* – dimensional representation of *kmers* or sequences in ℝ^*n*^. The architecture for `BioVec` and `Seq2vec` models are provided in Fig 3.

**Fig 3.**
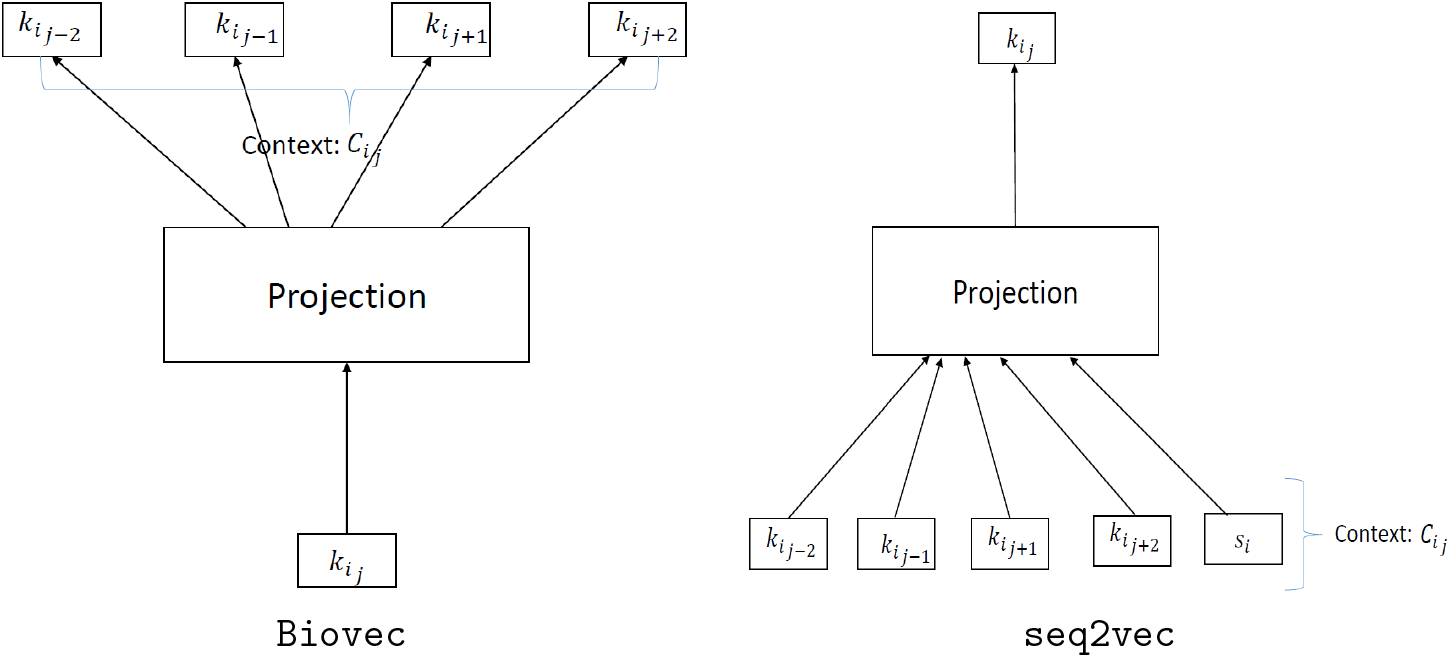
Architecture for `BioVec` and `Seq2Vec` models. *k_i_j__* represents *j^th^ k-mer* in the *i^th^* sequence, *s_i_*.

The training of these models involve a simple prediction mechanism over *samples,* consisting of *k-mer* and their neighbors (context) randomly picked from the corpus of sequences. For prediction over a *sample*, either the *kmer* is predicted from the context or vice-versa as depicted in Fig 3. In `BioVec` model, the *sample* consists of *k-mer* and its neighbours (context), whereas for `Seq2Vec`, the sample also contains the tag associated with the corresponding sentence. Mathematically, by iterating through the samples, the parameters of these models are learned such that negative log-likelihood probabilities of co-occurrence of *k-mers* is minimized. The cost function that is minimized in `BioVec` and `Seq2Vec` is given in Eq (1) and Eq (2) respectively.

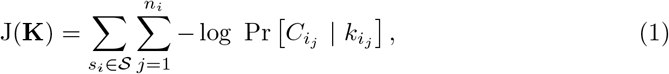

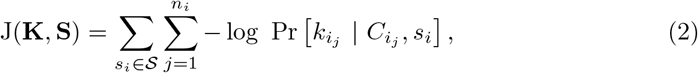

here *C_i_j__* (context) consist of the surrounding *k-mers* of *k_i_j__* and *s_i_* is the sequence tag.

Once the `BioVec` model is trained, the sequence embedding is computed by linearly combining the embeddings of its constituent *k-mers*. For `Seq2Vec`, the sequence embeddings are directly obtained by passing them through the trained model. This step is also known as an inference step. Note that these approaches are unsupervised and only need the sequence corpus for training.

The followup work by Kimothi et al. [35] presented supervised approaches for learning representations of protein sequences: `SuperVec` and `SuperVecX`. These supervised approaches use the sequences as well as their associated labels during the training of the models. Similar to `Word2Vec`, these models are trained via a prediction task. Training of `SuperVecX` involves prediction of the class label using all *k-mers* of given sequences.

In contrast to `SuperVecX`, the `SuperVec` model joins the two prediction units together to learn contextual and label information. The first unit follows the prediction of a *k-mer* given its context to capture contextual information. The second unit ensures the inclusion of label information in the training process by forcing the network to predict the tags of sequences that share the label with the considered *sample.* The joint cost function [35] of `SuperVec` is given as below,

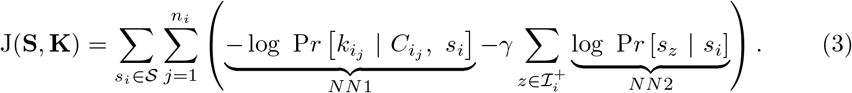

Here 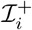 is the set of sequences that share label with *s_i_*. Note that NN1 and NN2 in Eq. (3) represents the two parts of the complete model, i.e. for learning contextual and label information respectively. As noted before, the `SuperVec` requires the labels of protein sequences for training purposes. Since for the PPI prediction problem, we also have the interaction network along with the sequences, we incorporate limited information from the network by exploiting it to obtain the sequence labels. We call this adaptation of `SuperVec` as `SuperVecNW`; the description is given below:

### SuperVecNW

In using `SuperVecNW` we make use of the available network information to label the sequences. The neighborhood of a sequence is looked up in the complete interaction network obtained from the training pairs of proteins. For a given sequence, the sequences connected to it in the PPI network are considered to come from the same class (interacting) and others from the different class (non-interacting).

For example, Fig 1 shows a PPI network that is derived from a given training pairs. Here, for *s*_3_, *s*_1_, *s*_7_ and *s*_8_ are from the same class (interacting) and all others are the different (non-interacting) class. The joint cost function for `SuperVecNW` is same as that for `SuperVec` as given in Eq (3), the difference is only in the process of deriving class labels for a sample; for the given example in Fig 1, for sample *s*_3_, 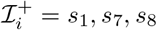 in Eq (3).

### 2.3 Proposed Framework

In the proposed framework, along with the sequence embedding techniques discussed above, we use the Random forest classifier for making predictions. Note that while the framework is generic and in principle any binary classifier can be used, we use the Random Forest Classifier [36] for our experiments due to its computational efficiency and convenience – only one hyper parameter needs to be tuned.

A Random Forest is a supervised classifier that uses an ensemble of decision trees to make predictions. These decision trees are constructed using bagging (a bootstrap aggregation technique [36]), and random feature selection. In the bagging method, each of the decision trees is trained using randomly drawn subsets of the training set. For random feature selection, the nodes in each decision tree are split based on a random selection of *m* features drawn from the n features available. Here, *m* <= *n*. For a test sample, the prediction is made based on voting (averaging) the prediction output from each decision tree. The presence of multiple trees in the random forest helps to avoid overfitting and limits the error due to bias. Also, there is virtually no hyper-parameter to tune except the number of trees.

In this paper, we represent each sequence as a 100-dimensional feature vector, and the pair of proteins as 200 dimensional vector. To use `SuperVecX`, we concatenate the sequences, with the result that the protein *pair* is represented by a *single* 100 dimensional feature vector. Note the much lower dimensionality of these representations compared to the other established feature vectors discussed in section 1.1. For interacting and non-interacting pairs, we use the class labels 1 and 0 respectively. The number of decision trees is fixed as 500, where each of the trees is built by selecting *m* ≪ 200 random features at each node. A test sample is then predicted based on voting of the decision trees.

The complete pipeline of the proposed framework is provided in Fig 2. Here block A shows the training stage of the representation learning model, A1; when trained, we denote it as A2. The A1 is trained using the sequences in training set 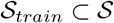. Note that 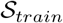 constitutes of all the unique sequences present in protein pairs in 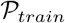. Block B is used for making predictions. It is operated in two phases, namely, the training and testing phase. In the training phase, the samples (protein pair) from 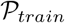 are passed through B1 to generate the embeddings of their constituent sequences. These embeddings are then concatenated to finally create the feature vector for the protein pairs, which are further used to train the binary classifier, B3. In the testing phase, similar to the training phase, the feature vector for any given test pair from 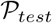 is generated by passing them through B1 and B2. This feature vector is finally given to the trained binary classifier for predicting if the test pair is interacting or not.

## 3 Results and Discussion

In this section, we present the results of our approach on the benchmark *human* and *yeast* PPI datasets provided by Park et al. [24]. These datasets were originally obtained from the protein interaction network analysis platform [37] and were further refined to contain only those sequences that share up to 40% identity. We also provide the results on another widely used benchmark *yeast* dataset that is provided by Guo et al. [15]; this dataset was originally obtained from DIP [38] and then refined such that majority of the protein pairs in it share less than 40% pairwise sequence identity.

### 3.1 Evaluation Scheme

For the experiments, we follow the evaluation scheme specifically proposed for the paired input problem by Park et al. [24]. In these problems the inference is made for a pair of objects (i.e. the PPI prediction) rather than for the single object (e.g., protein family prediction). Based on their experimental results, the authors in [24] showed that the performance of any method for the PPI prediction problem is affected by the presence of the individual proteins of the test sample in the training set, providing better results for the pairs that share protein(s) in the training data.

We cannot therefore expect a reliable assessment of generalisation if performance is evaluated on a test set dominated by samples which share components with the training data. Considering this issue, the authors suggested that the methods be evaluated on three different test classes, namely C1, C2, and C3. Here C1 constitutes the test samples of which both proteins are present in the training data, whereas in C2, only those test samples are considered that share one protein with the training data. Finally, the C3 test class is constructed from those test samples which share none of the constituent proteins with the training samples, making it the most difficult test class among the three.

Park et al. [24] also noted that the dataset previously used for cross-validation is close to the C1 type, but these pairs constitute only a modest fraction of known human PPIs. According to the HIPPIE database [44], the C1 type accounts for 19.2% of all interacting human protein pairs, whereas C2 and C3 constitute 49.2% and 31.6% of protein pairs respectively. Hence, in a typical cross-validation test set, the C1 class is more frequent than the population level, and performance estimates from cross-validation test sets can not be expected to generalize reliably. Therefore, apart from the evaluation on the cross-validation and C1 test classes, results obtained on C2 and C3 test classes should be taken into account before making any judgment about the generalization properties of the method.

### 3.2 Datasets

#### 3.2.1 Park and Marcotte datasets

We first evaluated our method on the Human and Yeast PPI datasets provided by Yungki Park and M.Marcotte [24]. The human PPI dataset contains 20,117 proteins and 24, 718 PPIs whereas the Yeast dataset contains 6, 806 proteins and 14, 938 PPIs. The experiments are conducted on the 40-fold train-test split for the CV, C1, C2, and C3 classes; for benchmarking the split files were also as provided by [24]. Results averaged over these splits are provided in Table 1 and 2.

**Table 1.** Comparison of the prediction performance between the proposed methods and other state of art methods on the Human dataset

| Methods | AUROC |  |  |  |
| --- | --- | --- | --- | --- |
|  | CV | C1 | C2 | C3 |
| M1 (Martin et al. [39]) | $0.81 \pm 0.01$ | $0.81 \pm 0.01$ | $0.61 \pm 0.01$ | $0.58 \pm 0.03$ |
| M2 (Vert et al. [40]) | $0.85 \pm 0.01$ | $0.85 \pm 0.01$ | $0.60 \pm 0.01$ | $0.58 \pm 0.02$ |
| M3 (Shen et al. [25]) | $0.63 \pm 0.01$ | $0.64 \pm 0.01$ | $0.55 \pm 0.01$ | $0.50 \pm 0.00$ |
| M4 (Guo et al. [15]) | $0.77 \pm 0.01$ | $0.77 \pm 0.01$ | $0.57 \pm 0.02$ | $0.53 \pm 0.02$ |
| M5 (Park et al. [24]) | $0.81 \pm 0.01$ | $0.81 \pm 0.01$ | $0.59 \pm 0.01$ | $0.54 \pm 0.03$ |
| M6 (Pitre et al. [41]) | $0.76 \pm 0.01$ | $0.77 \pm 0.01$ | $0.64 \pm 0.01$ | $0.59 \pm 0.02$ |
| M7 (Bellucci et al. [42]) | $0.56 \pm 0.01$ | $0.56 \pm 0.01$ | $0.53 \pm 0.01$ | $0.54 \pm 0.02$ |
| M8 (Ding et al. [43]) | $0.82 \pm 0.02$ | $0.82 \pm 0.01$ | $0.60 \pm 0.02$ | $0.57 \pm 0.02$ |
| **M9** (Seq2Vec) | $0.82 \pm 0.01$ | $0.82 \pm 0.01$ | $0.59 \pm 0.01$ | $0.55 \pm 0.02$ |
| **M10** (BioVec) | $0.77 \pm 0.02$ | $0.78 \pm 0.02$ | $0.59 \pm 0.01$ | $0.56 \pm 0.01$ |
| **M11** (SuperVecNW) | $0.82 \pm 0.04$ | $0.82 \pm 0.01$ | $0.58 \pm 0.01$ | $0.54 \pm 0.02$ |
| **M12** (SuperVecX) | $0.80 \pm 0.01$ | $0.80 \pm 0.01$ | $0.55 \pm 0.01$ | $0.52 \pm 0.02$ |
| SPRINT | NA | 0.82 | 0.66 | 0.61 |
| PIPR | $0.8 \pm 0.01$ | $0.8 \pm 0.01$ | $0.57 \pm 0.01$ | $0.52 \pm 0.01$ |

| Methods | AUPRC |  |  |  |
| --- | --- | --- | --- | --- |
|  | CV | C1 | C2 | C3 |
| M1 (Martin et al. [39]) | $0.82 \pm 0.01$ | $0.82 \pm 0.01$ | $0.60 \pm 0.01$ | $0.57 \pm 0.03$ |
| M2 (Vert et al. [40]) | $0.85 \pm 0.00$ | $0.85 \pm 0.01$ | $0.60 \pm 0.01$ | $0.56 \pm 0.03$ |
| M3 (Shen et al. [25]) | $0.67 \pm 0.01$ | $0.67 \pm 0.01$ | $0.57 \pm 0.02$ | $0.52 \pm 0.05$ |
| M4 (Guo et al. [15]) | $0.77 \pm 0.01$ | $0.77 \pm 0.01$ | $0.56 \pm 0.01$ | $0.53 \pm 0.02$ |
| M5 (Park et al. [24]) | $0.82 \pm 0.01$ | $0.82 \pm 0.01$ | $0.59 \pm 0.01$ | $0.54 \pm 0.02$ |
| M6 (Pitre et al. [41]) | $0.79 \pm 0.01$ | $0.79 \pm 0.01$ | $0.67 \pm 0.01$ | $0.59 \pm 0.02$ |
| M7 (Bellucci et al. [42]) | $0.56 \pm 0.01$ | $0.56 \pm 0.01$ | $0.53 \pm 0.01$ | $0.54 \pm 0.02$ |
| M8 (Ding et al. [43]) | $0.83 \pm 0.01$ | $0.83 \pm 0.01$ | $0.60 \pm 0.01$ | $0.56 \pm 0.02$ |
| **M9** (Seq2Vec) | $0.83 \pm 0.01$ | $0.83 \pm 0.01$ | $0.59 \pm 0.01$ | $0.55 \pm 0.02$ |
| **M10** (BioVec) | $0.78 \pm 0.01$ | $0.79 \pm 0.02$ | $0.59 \pm 0.01$ | $0.56 \pm 0.01$ |
| **M11** (SuperVecNW) | $0.84 \pm 0.02$ | $0.83 \pm 0.01$ | $0.59 \pm 0.01$ | $0.53 \pm 0.01$ |
| **M12** (SuperVecX) | $0.8 \pm 0.01$ | $0.8 \pm 0.01$ | $0.56 \pm 0.01$ | $0.53 \pm 0.01$ |
| SPRINT | NA | 0.86 | 0.70 | 0.63 |
| PIPR | $0.80 \pm 0.01$ | $0.80 \pm 0.01$ | $0.57 \pm 0.01$ | $0.51 \pm 0.01$ |

**Table 2.** Comparison of the prediction performance between the proposed methods and other state of art methods on the Yeast dataset

| Methods | AUROC |  |  |  |
| --- | --- | --- | --- | --- |
|  | CV | C1 | C2 | C3 |
| M1 (Martin et al. [39]) | $0.82 \pm 0.01$ | $0.82 \pm 0.01$ | $0.61 \pm 0.02$ | $0.58 \pm 0.03$ |
| M2 (Vert et al. [40]) | $0.83 \pm 0.01$ | $0.84 \pm 0.01$ | $0.60 \pm 0.02$ | $0.59 \pm 0.03$ |
| M3 (Shen et al. [25]) | $0.61 \pm 0.01$ | $0.61 \pm 0.01$ | $0.53 \pm 0.01$ | $0.50 \pm 0.01$ |
| M4 (Guo et al. [15]) | $0.76 \pm 0.02$ | $0.76 \pm 0.02$ | $0.57 \pm 0.02$ | $0.54 \pm 0.03$ |
| M5 (Park et al. [24]) | $0.80 \pm 0.02$ | $0.80 \pm 0.01$ | $0.58 \pm 0.01$ | $0.55 \pm 0.02$ |
| M6 (Pitre et al. [41]) | $0.75 \pm 0.02$ | $0.75 \pm 0.02$ | $0.59 \pm 0.04$ | $0.52 \pm 0.04$ |
| M7 (Bellucci et al. [42]) | $0.58 \pm 0.02$ | $0.58 \pm 0.01$ | $0.54 \pm 0.02$ | $0.52 \pm 0.03$ |
| M8 (Ding et al. [43]) | $0.82 \pm 0.02$ | $0.82 \pm 0.01$ | $0.62 \pm 0.02$ | $0.61 \pm 0.02$ |
| **M9** (Seq2Vec) | $0.81 \pm 0.02$ | $0.81 \pm 0.01$ | $0.59 \pm 0.02$ | $0.58 \pm 0.02$ |
| **M10** (BioVec) | $0.81 \pm 0.01$ | $0.81 \pm 0.01$ | $0.61 \pm 0.04$ | $0.61 \pm 0.02$ |
| **M11** (SuperVecNW) | $0.82 \pm 0.01$ | $0.8 \pm 0.01$ | $0.58 \pm 0.01$ | $0.54 \pm 0.01$ |
| **M12** (SuperVecX) | $0.81 \pm 0.01$ | $0.81 \pm 0.01$ | $0.55 \pm 0.01$ | $0.54 \pm 0.02$ |
| PIPR | $0.83 \pm 0.03$ | $0.84 \pm 0.01$ | $0.6 \pm 0.02$ | $0.56 \pm 0.03$ |

| Methods | AUPRC |  |  |  |
| --- | --- | --- | --- | --- |
|  | CV | C1 | C2 | C3 |
| M1 (Martin et al. [39]) | $0.83 \pm 0.02$ | $0.83 \pm 0.01$ | $0.62 \pm 0.02$ | $0.57 \pm 0.03$ |
| M2 (Vert et al. [40]) | $0.84 \pm 0.02$ | $0.84 \pm 0.01$ | $0.61 \pm 0.02$ | $0.58 \pm 0.03$ |
| M3 (Shen et al. [25]) | $0.65 \pm 0.02$ | $0.65 \pm 0.02$ | $0.56 \pm 0.03$ | $0.53 \pm 0.07$ |
| M4 (Guo et al. [15]) | $0.76 \pm 0.02$ | $0.76 \pm 0.02$ | $0.58 \pm 0.02$ | $0.54 \pm 0.03$ |
| M5 (Park et al. [24]) | $0.78 \pm 0.02$ | $0.78 \pm 0.01$ | $0.57 \pm 0.02$ | $0.54 \pm 0.02$ |
| M6 (Pitre et al. [41]) | $0.75 \pm 0.02$ | $0.76 \pm 0.02$ | $0.60 \pm 0.05$ | $0.47 \pm 0.07$ |
| M7 (Bellucci et al. [42]) | $0.60 \pm 0.02$ | $0.60 \pm 0.02$ | $0.55 \pm 0.025$ | $0.53 \pm 0.02$ |
| M8 (Ding et al. [43]) | $0.84 \pm 0.01$ | $0.84 \pm 0.01$ | $0.64 \pm 0.02$ | $0.62 \pm 0.02$ |
| **M9** (Seq2Vec) | $0.83 \pm 0.02$ | $0.83 \pm 0.01$ | $0.61 \pm 0.02$ | $0.59 \pm 0.02$ |
| **M10** (BioVec) | $0.82 \pm 0.01$ | $0.82 \pm 0.01$ | $0.62 \pm 0.03$ | $0.62 \pm 0.02$ |
| **M11** (SuperVecNW) | $0.84 \pm 0.02$ | $0.82 \pm 0.01$ | $0.59 \pm 0.01$ | $0.53 \pm 0.02$ |
| **M12** (SuperVecX) | $0.81 \pm 0.01$ | $0.56 \pm 0.01$ | $0.54 \pm 0.01$ | $0.54 \pm 0.01$ |
| PIPR | $0.83 \pm 0.03$ | $0.84 \pm 0.02$ | $0.6 \pm 0.02$ | $0.55 \pm 0.02$ |

#### 3.2.2 Guo dataset

Gua et al. [15] provide PPI datasets for several species. These datasets are balanced and contain an equal number of positive and negative samples. In this paper, we use the yeast dataset used as a benchmark in many of the earlier studies [28, 31]. In this dataset, there are 2450 unique proteins, and a total of 11188 protein pairs, where half of these pairs are interacting (positive pair) and rest are non-interacting (negative pair). For negative pairs, the proteins that have no evidence of interactions are randomly selected, that are further filtered based on their sub-cellular location.

### 3.3 Evaluation Measurements

Park et al. [24] use AUROC – the area under the Receiver Operating Characteristic curve – and the mean of the AUPRC – the area under the Precision-Recall curve – to benchmark the performance of previous studies. The ROC is a graph showing the relationship between the False positive rate (FPR) and the True positive rate (TPR), whereas the PRC is the graph showing precision and recall values calculated at different thresholds. The AUROC and AUPRC summarize the ROC and PR curves, respectively. The AUROC value indicates the ability of the classifier to distinguish between two classes, whereas AUPRC shows the trade-off achieved between precision and recall. To compare with these benchmark methods, we also compute the AUROC and AUPRC values (averaged for 40 randomly selected train-test splits) to evaluate our approaches for the *Park datasets.* The benchmark methods that use the *Guo dataset* for performance evaluation use accuracy, precision, sensitivity, specificity, F1-score, and MCC as evaluation metrics. To compare with these benchmark methods, we evaluate our proposed approaches using these same metrics.

The evaluation metrics used are defined as follows:

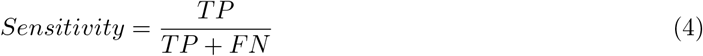

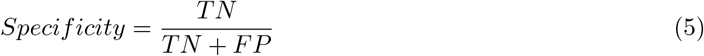

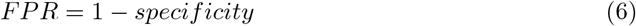

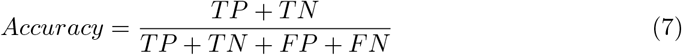

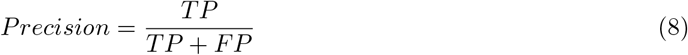

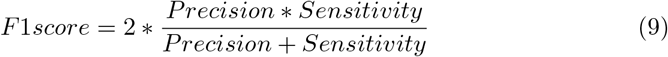

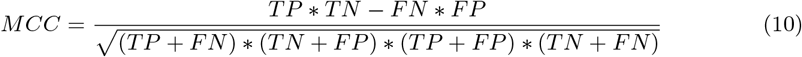

Here the True Positive (TP) and True Negative (TN) counts record the number of correctly predicted interacting and noninteracting protein pairs. Similarly, the False Negative (FN) and False Positive (FP) counts reflect the number of incorrectly predicted non-interacting and interacting pairs, respectively.

### 3.4 Experiments and Results

#### 3.4.1 Experiments on *Park’s dataset*

In these experiments, we compare the results of the proposed framework with benchmark methods (M1-M8) that rely on physico-chemical properties for encoding the protein sequences. Here, M1 and M2 are signature-product based methods proposed by Martin et al. [39]. M1 uses the signature-product directly, whereas M2 [40] uses metric learning on the signatures to compute the feature vector for the protein pair. M3 [25] is the Conjoint triad method; in this method, the feature vector of a protein sequence is based on the frequency of conjoint triads. M4 and M5 both use the AC method proposed by Guo et al. [15] for feature construction, but they differ in the classification algorithm, with M4 using the SVM whereas M5 employs a random forest. The M7 method was developed initially for protein-RNA interaction prediction and is based on maximizing the scores computed for interacting protein pairs vis-a-vis non-interacting protein pairs within the vector space. M8, proposed by Ding et al. [43], calculates mutual information-based features based on the functional categories of amino acids. For predictions, M1-M4 use the SVM while M5-M8 use a random forest classifier. All of these methods (M1-M8), except M6 (PIPE), are machine learning-based approaches. PIPE is a sequence similarity-based approach that relies on the shared local similarity between known interacting protein pairs and the test pair. In the comparison for the human dataset, we also report another sequence similarity-based approach, SPRINT [45]. In contrast to the machine learning-based approaches, these methods use only positive (interacting) pairs for making predictions and hence do not require the negative (non-interacting) examples.

Our approach is a generic machine learning based framework in which we can accommodate different representation learning methods for generating the feature vectors. We report the results for four different representation learning techniques – `Seq2Vec` (M9), `BioVec` (M10), `SuperVecNW` (M11) and `SuperVecX` (M12) – on both the *human* and *yeast* datasets.

##### Results

Table 1 and Table 2 summarize the performance of each method on the *human* and *yeast* datasets respectively. We report the mean AUROC and AUPRC and their corresponding standard deviations calculated over forty randomly selected train-test splits.

On the human dataset, for the CV and C1 test set, the performance of our methods (M9, M11, and M12) is better than M1-M7 except for M2 and gives comparable performance to M8 and SPRINT. When compared to the conjoint triad (M3) method, we achieve an improvement of 18%.

For C2, the averaged AUROC value of all machine learning-based methods ranges between 0.55 — 0.61 whereas PIPE (M6) and SPRINT provide comparatively better results, i.e., 0.64 and 0.66 respectively. For C3, the values of averaged AUROC for all methods range from 0.55 — 0.61. Note that for a classifier that randomly predicts interacting and non-interacting pairs, the AUROC is expected to be 0.5 and hence none of these methods can be considered to perform very well for C3. This leaves considerable scope for improvement in the feature construction methods. The AUPRC values obtained for all methods follow the same trend as AUROC values. Overall, SPRINT provides the best results for the C2 and C3 test class.

The results obtained for all methods on the *yeast* dataset are in general consistent with the results obtained for the *human* dataset, except for PIPE, which deteriorates for C2 and C3. SPRINT results are not available for the *yeast* dataset.

Comparing the performance of different RepL techniques i.e. `Seq2Vec` (M9), `BioVec` (M10), `SuperVecNW` (M11) and `SuperVecX` (M12) on the *human* dataset, we observe that for CV and C1, M9, M11 and M12 provide slightly better result than M10, whereas for C2 and C3 they provide comparable values. For the yeast data set, M9 and M10 provide comparable results. This difference in performance when compared to the human dataset might be due to the larger number of *human* protein interactions available for training as compared to the *yeast* dataset.

In summary, our proposed framework (tested with different RepL approaches) outperforms most of the standard methods for CV and C1, while giving comparable results for C2 and C3 in most cases. We also compare the proposed framework with a recent deep learning based architecture – PIPR, as discussed below.

##### Comparison with Deep Learning approach – PIPR

Some recent work has demonstrated the applicability of deep learning based approaches for the PPI prediction task. All of these approaches are primarily based on the Convolutional Neural Network and have a deep architecture. The most recent of these approaches, PIPR [33], provides an end-to-end deep GRU (gated recurrent unit) based architecture for PPI prediction. The results reported [33] show an improvement in classification results (refer Table 3) compared to other benchmark methods – including other deep learning based approaches.

**Table 3.** Evaluation of PPI prediction on the Yeast dataset (Guo’s) based on 5 fold cross validation. Mean and standard deviation is reported in the table

| Methods | Accuracy (%) | Precision (%) | Sensitivity (%) | Specificity (%) | F1-score (%) | MCC (%) |
| --- | --- | --- | --- | --- | --- | --- |
| SVM-AC | 87.35 ± 1.38 | 87.82 ± 4.84 | 87.30 ± 5.23 | 87.41 ± 6.33 | 87.34 ± 1.33 | 75.09 ± 2.51 |
| kNN-CTD | 86.15 ± 1.17 | 90.24 ± 1.34 | 81.03 ± 1.74 | NA | 85.39 ± 1.51 | NA |
| EELM-PCA | 86.99 ± 0.29 | 87.59 ± 0.32 | 86.15 ± 0.43 | NA | 86.86 ± 0.37 | 77.36 ± 0.44 |
| SVM-MCD | 91.36 ± 0.4 | 91.94 ± 0.69 | 90.67 ± 0.77 | NA | 91.3 ± 0.73 | 84.21 ± 0.66 |
| MLP | 94.43 ± 0.3 | 96.65 ± 0.59 | 92.06 ± 0.36 | NA | 94.3 ± 0.45 | 88.97 ± 0.62 |
| RF-LPQ | 93.92 ± 0.36 | 96.45 ± 0.45 | 91.10 ± 0.31 | NA | 93.7 ± 0.37 | 88.56 ± 0.63 |
| SAE | 67.17 ± 0.62 | 66.90 ± 1.42 | 68.06 ± 2.50 | 66.30 ± 2.27 | 67.44 ± 1.08 | 34.39 ± 1.25 |
| DNN-PPI | 76.61 ± 0.51 | 75.1 ± 0.66 | 79.63 ± 1.34 | 73.59 ± 1.28 | 77.29 ± 0.66 | 53.32 ± 1.05 |
| DPPI | 94.55 | 96.68 | 92.24 | NA | 94.41 | NA |
| SRGRU | 93.77 ± 0.84 | 94.60 ± 0.64 | 92.85 ± 1.58 | 94.69 ± 0.81 | 93.71 ± 0.85 | 87.56 ± 1.67 |
| SCNN | 95.03 ± 0.47 | 95.51 ± 0.77 | 94.51 ± 1.27 | 95.55 ± 0.77 | 95.00 ± 0.50 | 90.08 ± 0.93 |
| PIPR | 97.09 ± 0.24 | 97.00 ± 0.65 | 97.17 ± 0.44 | 97.00 ± 0.67 | 97.09 ± 0.23 | 94.17 ± 0.48 |
| **M9** (Seq2Vec) | 94.24 ± 0.48 | 97.35 ± 0.55 | 90.97 ± 0.91 | 97.52 ± 0.53 | 94.05 ± 0.51 | 88.68 ± 0.93 |
| **M10** (BioVec) | 94.18 ± 0.30 | 97.16 ± 0.67 | 91.03 ± 0.67 | 97.34 ± 0.65 | 93.99 ± 0.31 | 88.55 ± 0.61 |

These results are computed following a 5-fold cross validation test on Guo’s yeast dataset. As discussed before, the cross-validation results may not reflect the ability of the method to generalize over the complete dataset. We thus evaluate the performance of PIPR on *Marcotte’s* dataset for C1, C2, C3 test classes along with the cross validation setting. The results for *yeast* dataset show improvement (2-3%) for CV and C1; for C2 the results are comparable but for C3 the figure is some 2-4% below our approaches. For the human dataset, its performance is generally 2-4% below our methods. These results show that like the other techniques PIPR does not perform well for C2 and C3 test sets. For the human dataset, it is not the best among the machine learning based methods.

#### 3.4.2 Experiment with Guo’s dataset

In these experiments, we compare the results of our methods (M9 and M10) on Guo’s yeast dataset with baseline approaches SVM-AC [15], kNN-CTD [17], EELM-PCA [46], SVM-MCD [27], MLP [47], RF-LPQ [28], SAE [6], DNN-PPI [32], DPPI [31] and SRGRU, SCNN and PIPR from [33]. The results for the baseline approaches are taken from [33].

##### Results

As shown in Table 3, M9 and M10 outperform many approaches (SVM-AC, kNN-CTD, EELM-PCA, SVM-MCD, SAE, DNN-PPI, SRGRU) and perform at par with DPPI and SCNN. The only method that outperforms our approaches is PIPR with an improvement of approx 2.5% in accuracy. Note that PIPR is a deep end-to-end method whereas our methods comprise a simple feature extraction method with a binary classifier. Also, it is important to mention that these results are reported in the cross-validation setting and hence should not be considered to generalize for the complete dataset. It is appropriate to evaluate these further on the C2 and C3 test classes. As discussed before we evaluated PIPR on Marcotte’s benchmark dataset and procedure (reported in Table ?? and ??) and find that even deep learning methods like PIPR do not perform well for C2 and C3.

### 3.5 Discussion

Predicting PPIs is a challenging problem. In recent years, computational methods have been a valuable alternative to experimental methods owing to their speed and relatively low false-positive rates. Over this period, sequence-based techniques have evolved, and better performance – as much as 18% compared to the earliest methods – has been achieved. Sequence-based methods mainly differ in the way the feature vector is constructed from the sequences. Most of the available methods rely on known physico-chemical properties of the protein. Such properties might not cover the interaction information completely and hence may not be useful in distinguishing interacting from non-interacting pairs. The availability of sequence data and the improvement in automatic extraction of features in text processing have motivated researchers to learn feature vectors directly from the sequences. This work establishes the fact that representation learning (RepL) based approaches can produce sequence embeddings that are useful for PPI prediction, providing better results than the physico-chemical property feature vectors without the need for extensive domain knowledge.

While sequence-based methods have evolved and have been shown to provide a high level of accuracy for PPI prediction, we need to be careful while making a judgment about their ability to generalize. Most of these methods give high cross-validation accuracy, which may not be a reflection on their generalization properties [24]. Therefore, we need to also focus on the results of the C2 and C3 test classes.

In our experiments, we found that all methods – including the state-of-art deep learning method PIPR – show good cross-validation accuracy, but when evaluated for C2 and C3 test class, their performance markedly deteriorates. Based on these observations, it can be inferred that capturing the interaction information from a pair of proteins in a feature vector remains a challenging task. While our methods offer much more convenient construction of feature vectors than some existing alternatives, and performance remains comparable, some insight is lacking and there is a good deal of scope to improve our models for generating feature vectors from raw sequences.

## 4 Conclusions

In this paper, we proposed a framework that uses the embeddings learned through `word2vec` based models for the PPI prediction task. Such embeddings do not require any prior biological knowledge and are learned directly from the sequence data. Moreover, these embeddings offer a far lower dimensional representation than those based on physico-chemical properties of the proteins and their constituent amino acids.

The experimental results obtained on both Human and Yeast datasets confirm that better sequence embeddings for PPI prediction tasks can be generated without necessarily relying on biological prior knowledge. We observe that most of the sequence-based methods perform well when evaluated on CV and C1 but not on the C2 and C3 test classes. The inability of sequence-based methods to perform better for C2 and C3 test classes presents an attractive if challenging opportunity to learn feature vectors that better represent the interaction between two proteins. Perhaps some limited use of prior knowledge in a supervised learning framework may provide the answer – maintaining most of the convenience of our approach while offering improved performance on the more challenging datasets.

## References

1. De Las Rivas J, Fontanillo C. Protein–protein interactions essentials: key concepts to building and analyzing interactome networks. PLoS computational biology. 2010;6(6):e1000807.

2. Wang RS, Wang Y, Wu LY, Zhang XS, Chen L. Analysis on multi-domain cooperation for predicting protein–protein interactions. BMC bioinformatics. 2007;8(1):391.

3. Koegl M, Uetz P. Improving yeast two-hybrid screening systems. Briefings in Functional Genomics and Proteomics. 2007;6(4):302–312.

4. Nagamine N, Sakakibara Y. Statistical prediction of protein–chemical interactions based on chemical structure and mass spectrometry data. Bioinformatics. 2007;23(15):2004–2012.

5. Rüetschi U, Rosén Å, Karlsson G, Zetterberg H, Rymo L, Hagberg H, et al. Proteomic analysis using protein chips to detect biomarkers in cervical and amniotic fluid in women with intra-amniotic inflammation. Journal of proteome research. 2005;4(6):2236–2242.

6. Sun T, Zhou B, Lai L, Pei J. Sequence-based prediction of protein protein interaction using a deep-learning algorithm. BMC bioinformatics. 2017;18(1):277.

7. Han DS, Kim HS, Jang WH, Lee SD, Suh JK. PreSPI: a domain combination based prediction system for protein–protein interaction. Nucleic acids research. 2004;32(21):6312–6320.

8. Wan KK, Park J, Suh JK. Large scale statistical prediction of protein–protein interaction by potentially interacting domain (PID) pair. Genome Informatics. 2002;13:42–50.

9. Sprinzak E, Margalit H. Correlated sequence-signatures as markers of protein-protein interaction. Journal of molecular biology. 2001;311(4):681–692.

10. Huang TW, Tien AC, Huang WS, Lee YCG, Peng CL, Tseng HH, et al. POINT: a database for the prediction of protein–protein interactions based on the orthologous interactome. Bioinformatics. 2004;20(17):3273–3276.

11. Espadaler J, Romero-Isart O, Jackson RM, Oliva B. Prediction of protein–protein interactions using distant conservation of sequence patterns and structure relationships. Bioinformatics. 2005;21(16):3360–3368.

12. Ogmen U, Keskin O, Aytuna AS, Nussinov R, Gursoy A. PRISM: protein interactions by structural matching. Nucleic acids research. 2005;33(suppl_2):W331–W336.

13. Aloy P, Russell RB. InterPreTS: Protein Inter action Pre diction through Tertiary S tructure. Bioinformatics. 2003;19(1):161–162.

14. Aloy P, Russell RB. Interrogating protein interaction networks through structural biology. Proceedings of the National Academy of Sciences. 2002;99(9):5896–5901.

15. Guo Y, Yu L, Wen Z, Li M. Using support vector machine combined with auto covariance to predict protein-protein interactions from protein sequences. Nucleic acids research. 2008;36(9):3025–3030.

16. Zhou YZ, Gao Y, Zheng YY. Prediction of protein-protein interactions using local description of amino acid sequence. In: Advances in Computer Science and Education Applications. Springer; 2011. p. 254–262.

17. Yang L, Xia JF, Gui J. Prediction of protein-protein interactions from protein sequence using local descriptors. Protein and Peptide Letters. 2010;17(9):1085–1090.

18. Asgari E, Mofrad MRK. Continuous Distributed Representation of Biological Sequences for Deep Proteomics and Genomics. PLOS ONE. 2015;10(11):e0141287. doi:10.1371/journal.pone.0141287.

19. Mikolov T, Corrado G, Chen K, Dean J. Efficient Estimation of Word Representations in Vector Space. Proceedings of the International Conference on Learning Representations (ICLR 2013). 2013; p. 1–12. doi:10.1162/153244303322533223.

20. Ng P. dna2vec: Consistent vector representations of variable-length k-mers. arXiv preprint arXiv:170106279. 2017;.

21. Yang KK, Wu Z, Bedbrook CN, Arnold FH. Learned protein embeddings for machine learning. Bioinformatics. 2018;1:7.

22. Schwartz AS, Hannum GJ, Dwiel ZR, Smoot ME, Grant AR, Knight JM, et al. Deep semantic protein representation for annotation, discovery, and engineering. BioRxiv. 2018; p. 365965.

23. Asgari E, McHardy AC, Mofrad MR. Probabilistic variable-length segmentation of protein sequences for discriminative motif discovery (DiMotif) and sequence embedding (ProtVecX). Scientific reports. 2019;9(1):3577.

24. Park Y, Marcotte EM. Flaws in evaluation schemes for pair-input computational predictions. Nature methods. 2012;9(12):1134.

25. Shen J, Zhang J, Luo X, Zhu W, Yu K, Chen K, et al. Predicting protein–protein interactions based only on sequences information. Proceedings of the National Academy of Sciences. 2007;104(11):4337–4341.

26. You ZH, Chan KC, Hu P. Predicting protein–protein interactions from primary protein sequences using a novel multi-scale local feature representation scheme and the random forest. PLoS One. 2015;10(5):e0125811.

27. You ZH, Zhu L, Zheng CH, Yu HJ, Deng SP, Ji Z. Prediction of protein-protein interactions from amino acid sequences using a novel multi-scale continuous and discontinuous feature set. In: BMC bioinformatics. vol. 15. BioMed Central; 2014. p. S9.

28. Wong L, You ZH, Li S, Huang YA, Liu G. Detection of protein-protein interactions from amino acid sequences using a rotation forest model with a novel pr-lpq descriptor. In: International Conference on Intelligent Computing. Springer; 2015. p. 713–720.

29. Huang YA, You ZH, Gao X, Wong L, Wang L. Using weighted sparse representation model combined with discrete cosine transformation to predict protein-protein interactions from protein sequence. BioMed research international. 2015;2015.

30. Henikoff S, Henikoff JG. Amino acid substitution matrices from protein blocks. Proceedings of the National Academy of Sciences. 1992;89(22):10915–10919.

31. Hashemifar S, Neyshabur B, Khan AA, Xu J. Predicting protein–protein interactions through sequence-based deep learning. Bioinformatics. 2018;34(17):i802–i810.

32. Li H, Gong XJ, Yu H, Zhou C. Deep Neural Network Based Predictions of Protein Interactions Using Primary Sequences. Molecules. 2018;23(8):1923.

33. Chen M, Ju CJT, Zhou G, Chen X, Zhang T, Chang KW, et al. Multifaceted protein-protein interaction prediction based on Siamese residual RCNN. Bioinformatics. 2019;35(14):i305–i314.

34. Kimothi D, Soni A, Biyani P, Hogan JM. Distributed Representations for Biological Sequence Analysis. arXiv preprint arXiv:160805949. 2016;.

35. Kimothi D, Biyani P, Hogan JM, Soni A, Kelly W. Learning supervised embeddings for large scale sequence comparisons. BioRxiv. 2019; p. 620153.

36. Breiman L. Random forests. Machine learning. 2001;45(1):5–32.

37. Wu J, Vallenius T, Ovaska K, Westermarck J, Mäkelä TP, Hautaniemi S. Integrated network analysis platform for protein-protein interactions. Nature methods. 2009;6(1):75.

38. Xenarios I, Salwinski L, Duan XJ, Higney P, Kim SM, Eisenberg D. DIP, the Database of Interacting Proteins: a research tool for studying cellular networks of protein interactions. Nucleic acids research. 2002;30(1):303–305.

39. Martin S, Roe D, Faulon JL. Predicting protein–protein interactions using signature products. Bioinformatics. 2004;21(2):218–226.

40. Vert JP, Qiu J, Noble WS. A new pairwise kernel for biological network inference with support vector machines. In: BMC bioinformatics. vol. 8. BioMed Central; 2007. p. S8.

41. Pitre S, North C, Alamgir M, Jessulat M, Chan A, Luo X, et al. Global investigation of protein–protein interactions in yeast Saccharomyces cerevisiae using re-occurring short polypeptide sequences. Nucleic acids research. 2008;36(13):4286–4294.

42. Bellucci M, Agostini F, Masin M, Tartaglia GG. Predicting protein associations with long noncoding RNAs. Nature methods. 2011;8(6):444.

43. Ding Y, Tang J, Guo F. Predicting protein–protein interactions via multivariate mutual information of protein sequences. BMC bioinformatics. 2016;17(1):398.

44. Schaefer MH, Fontaine JF, Vinayagam A, Porras P, Wanker EE, Andrade-Navarro MA. HIPPIE: Integrating protein interaction networks with experiment based quality scores. PloS one. 2012;7(2):e31826.

45. Li Y, Ilie L. SPRINT: ultrafast protein-protein interaction prediction of the entire human interactome. BMC bioinformatics. 2017;18(1):485.

46. You ZH, Lei YK, Zhu L, Xia J, Wang B. Prediction of protein-protein interactions from amino acid sequences with ensemble extreme learning machines and principal component analysis. In: BMC bioinformatics. vol. 14. BioMed Central; 2013. p. S10.

47. Du X, Sun S, Hu C, Yao Y, Yan Y, Zhang Y. DeepPPI: boosting prediction of protein–protein interactions with deep neural networks. Journal of chemical information and modeling. 2017;57(6):1499–1510.

